# Estimating disease spread using structured coalescent and birth-death models: A quantitative comparison

**DOI:** 10.1101/2020.11.30.403741

**Authors:** Sophie Seidel, Tanja Stadler, Timothy G. Vaughan

## Abstract

Understanding how disease transmission occurs between subpopulations is critically important for guiding disease control efforts irrespective of whether the subpopulations represent geographically separated people, age or risk groups. The structured coalescent (SC) and the multitype birth-death (MBD) model can both be used to infer migration rates between subpopulations from phylogenies reconstructed from pathogen genetic sequences. However, the two classes of phylodynamic methods rely on different assumptions.

Here, we report on a simulation study which compares inferences made using these models for a variety of migration rates in both endemic diseases and epidemic outbreaks. For the epidemic outbreak, we found that the MBD recovers the true migration rates better than the SC regardless of migration rate. We hypothesize that the inaccurate SC estimates stem from the its assumption of a constant population size. For the endemic scenario, our analysis shows that both models obtain a similar coverage of the migration rates, while the SC provides slightly narrower posterior intervals. Irrespective of the scenario, both models estimate the root location with similar coverage.

Our study provides concrete modelling advice for infectious disease analysts. For endemic disease either model can be used, while for epidemic outbreaks the MBD should be the model of choice. Additionally, our study reveals the need to develop the SC further such that varying population sizes can easily be taken into account.

**Author summary:** Controlling an infectious disease requires us to quantify and understand how it spreads through pools of susceptible individuals, defined by their belonging to different geographical regions, age or risk groups. Rates of pathogen movement between these pools can be inferred from pathogen phylogenies which are themselves reconstructed from pathogen genetic sequences collected from infected individuals. Two popular foundations for such models are the multitype birth-death model and the structured coalescent.

Although these models fulfill the same purpose, they differ in their assumptions and can, hence, produce contrasting results. To assess the appropriateness of the models in different situations, we performed a simulation study. We find that, for endemic diseases, both models are able to estimate the migration parameters reliably. For epidemic outbreaks, however, the multitype birth-death model obtains better estimates of the migration rates. We hypothesize that the structured coalescent’s inaccurate estimates for the epidemic scenario arise because it assumes a constant number of infected individuals through time.

## Introduction

Recent years have seen a rise in both the use of phylodynamic models and their complexity. Early models focused on inferring important population dynamic parameters, e.g. the reproductive number, from pathogen genetic data for a panmictic population [1]. However, a population is often structured into subpopulations, e.g. an infected individual is more likely to transmit to other individuals that are in the same region, risk group or share another attribute [2]. More recent models [3–7] can estimate the population dynamics within subpopulations and the migration rates between them. For concreteness, we will regard subpopulations as consisting of infected individuals in distinct geographical regions in the remainder of the paper.

In a Bayesian phylodynamic analysis of an infectious disease dataset, the input data is an alignment of pathogen sequences collected from infected individuals. To reconstruct the phylogenetic tree together with the DNA substitution parameters, and the past population dynamics, a variety of methods are available [8–10]. An important component of such an analysis is the *tree prior*. When formulated in terms of a generative model, this allows researchers to characterise the posterior distribution of the population dynamic parameters and, for structured populations, the migration rates between the subpopulations.

Two popular models for the tree prior in structured populations are the multitype birth-death (MBD) [11–13] and the structured coalescent (SC) [14–16]. Both describe a stochastic process, that generates a distribution of phylogenetic trees for a given parameter set. Nevertheless, the MBD and the SC differ substantially for a number of reasons. Firstly, the MBD explicitly models how the sampled sequences were collected from the total population. On the one hand, this allows the MBD to use additional signal stemming from the sampling times [17]. On the other hand, its results can be biased, if the sampling process is unknown and wrongly specified in the analysis. The SC generally conditions on the number and timing of the samples, even though exceptions exist [17–19]. Hence, commonly used implementations (such as the SC in BEAST 2) cannot make use of this information. Secondly, the two models treat population size changes differently. For the original SC, the population size has to stay constant in each region. If additional predictor data is available, the population dynamic parameters are allowed to vary through time in a piecewise constant fashion [20]. In contrast, the MBD allows the population size to change stochastically through time. How important are population size fluctuations in real world infectious disease analyses? If a disease is endemic in a population, the number of infected remains approximately at the same baseline level. But during an epidemic outbreak, the population grows exponentially. At the beginning of an outbreak, stochastic variation dominates the dynamics [21].

Given the models’ different strengths and weaknesses it is not necessarily straightforward to identify which one should be used in a given application. For unstructured populations, coalescent and birth-death models have been compared in simulation studies [17,22]. However, an analysis of the potential biases in the more complex case of structured populations is lacking. In this paper, we investigate the performance of the SC, as implemented in MultiTypeTree [3] (in particular assuming constant population sizes and conditioning on sampling), and the MBD, as implemented in BDMM [5], in two practically relevant situations: an epidemic outbreak and an endemic situation. For the epidemic analysis, we simulate trees under the MBD with exponential growth and incomplete sampling. In the endemic case, we generate trees under the SC with constant population size. Then, for each scenario, we use both models to infer the migration rates and root locations.

## Results

To compare the structured coalescent (SC) and the multitype birth-death (MBD) models, we analysed their performance on simulated data mimicking endemic and epidemic, infectious diseases. In the endemic scenario, we simulated phylogenetic trees using the SC with a constant population size (Fig 1, left panel). For the epidemic outbreak, we used the MBD with exponential growth (Fig 1, right panel), which captures the stochastic population size changes. For each disease scenario, we simulated 100 trees for varying migration rates (fast, medium, slow; details in Materials and methods) between two locations *i,j* ∈ {*r, b*}.

**Fig 1.**
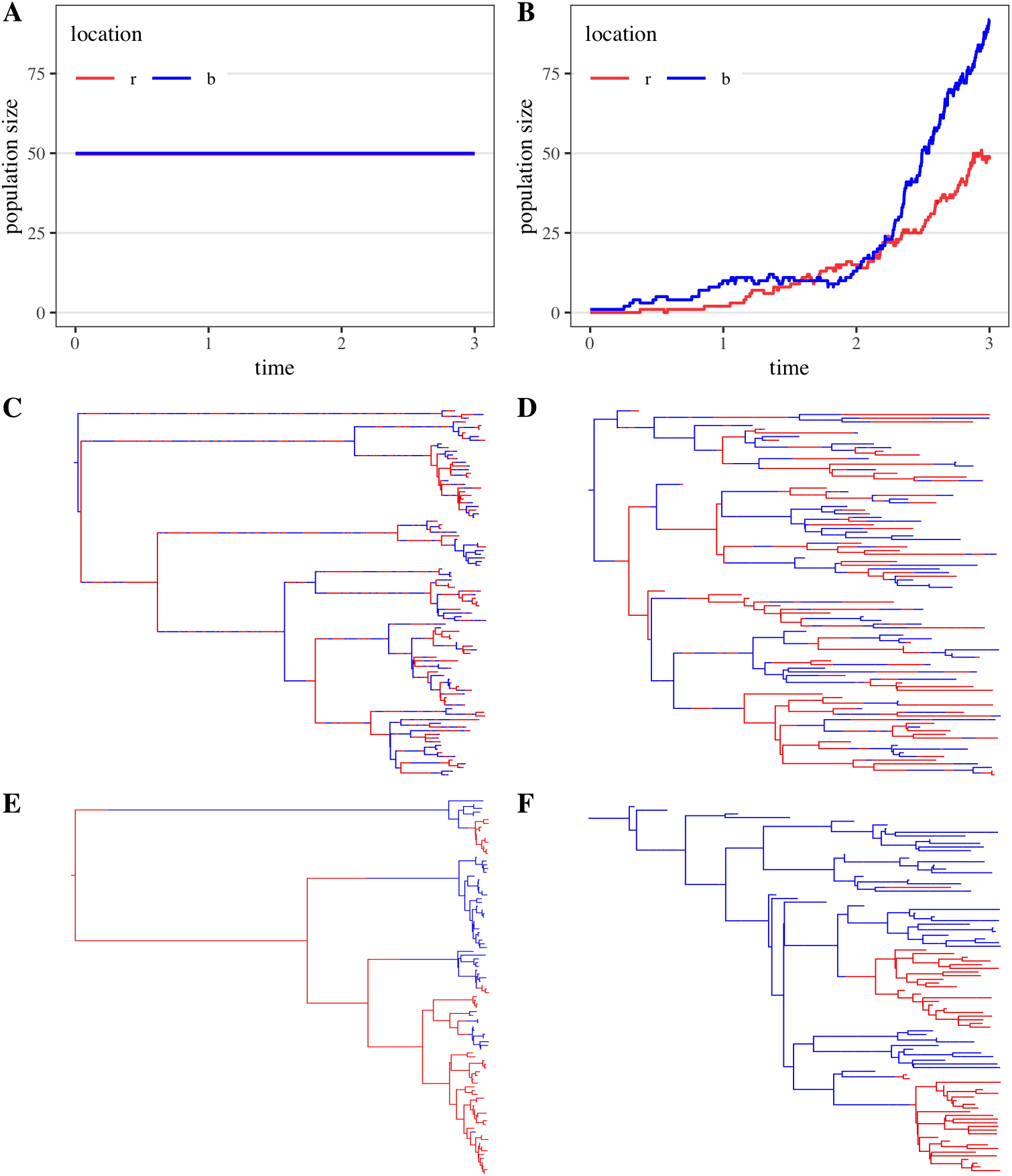
The endemic and epidemic scenario. We contrast the endemic (left) with the epidemic scenario (right). (**A**) The population size in each location {*r, b*} stays deterministically constant through time. (**B**) The population size in each location varies stochastically but grows exponentially in expectation (given the population does not go extinct). (**C-F**) Simulated, coloured trees. Branch colour indicates the location of an individual. (**C, E**) SC trees with samples from both locations. Note, that the branches close to the root are long compared to the branches close to the leaves of the tree. We show one tree for the fast (**C**) and for the slow (**E**) migration scenario. (**D, F**) MBD trees with samples from both locations. Here, the branches close to the leaves are long compared to the branches in root proximity. The trees were generated under fast (**D**) and slow (**F**) migration.

Then, we applied the SC and the MBD to infer the posterior distribution over the migration rates. We evaluate their performance using three metrics. Firstly, the coverage assesses how often the true migration rate is contained within the 95% highest posterior density (HPD) interval of the posterior. As a method can recover a parameter arbitrarily well by having wider HPD intervals, we also compare the relative HPD width. Additionally, we determine the relative root mean square error (RMSE) of each method. Here, a relative metric is attained by dividing the original metric by the true migration rate. For every migration case, each method infers 100 posterior distributions (one for every simulated tree). Therefore, we report the mean metrics and their standard deviation per parameter case and model.

### Migration rate inference in the endemic scenario

The results for the endemic scenario are summarised in Fig 2, left column. For fast migration, the MBD recovers the true migration rate 99% of the time. The SC achieves a slightly lower mean coverage of 95%. Further, the MBD yields an HPD width of around 4 which is 1.5 times narrower than the SC result (≥ 6). Similarly, the MBD is more accurate as indicated by its RMSE of ~ 0.5 which is 5 times lower compared to the RMSE of the SC. For medium and, especially, slow migration, the models perform comparably.

**Fig 2.**
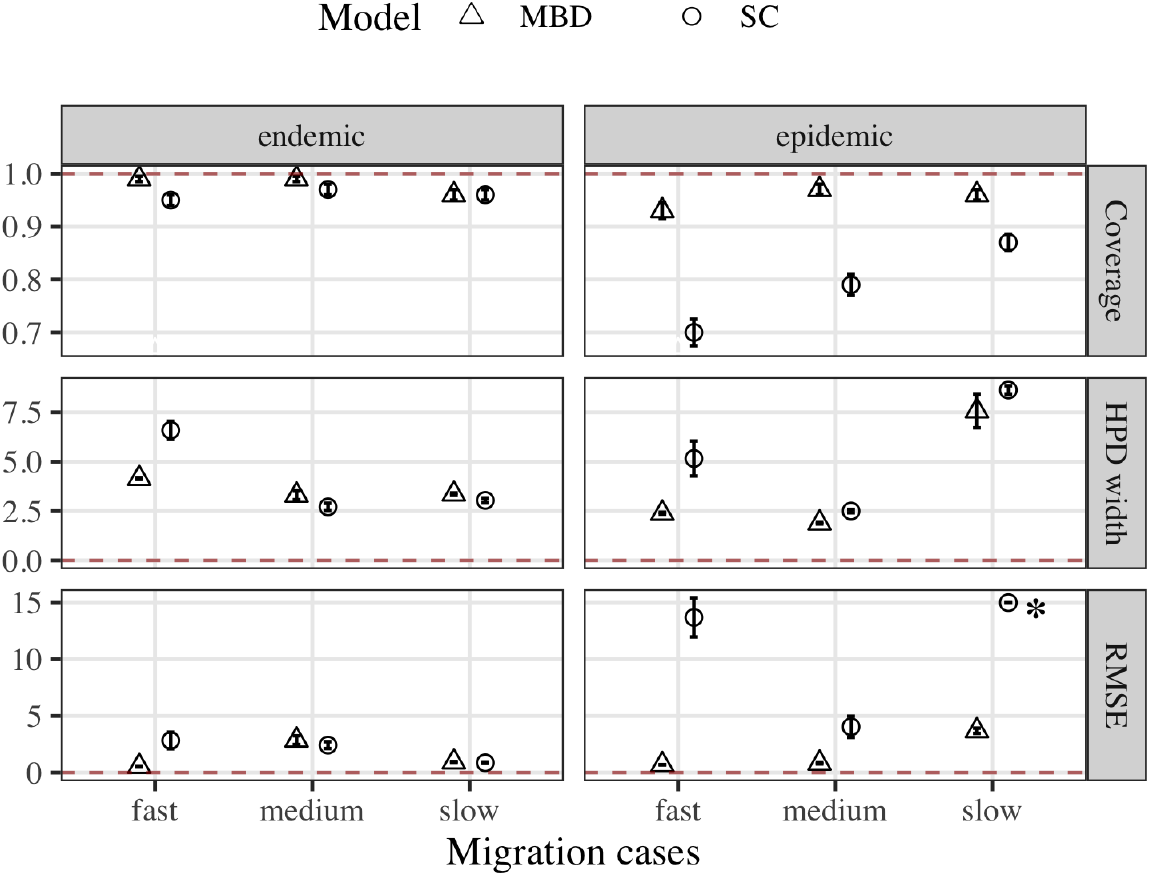
Summary statistics for migration rate inference. We report the summary statistics for the endemic (left) and the epidemic scenario with low sampling (right). For every migration case (fast, medium, slow), we summarise the inference results of the MBD (triangle) and the SC (circle) by reporting the coverage, relative HPD width and relative RMSE for *m_rb_*. Error bars represent the standard deviation of 1000 bootstrap samples around the mean. The red dashed line displays the optimal result for each metric. The SC’s RMSE in the epidemic scenario for slow migration (see *) was cut off for better visibility. Its value is 79.5 ± 24. (Results for *m_br_* can be found in S1 Fig)

It is curious that for fast migration, the model used for simulation (SC) does not perform as well as the MBD. Therefore, we investigated whether the prior distribution on the migration rate affected the outcome. To provide each model with the same information we had used the identical expontial prior with rate one. However, in the BEAST 2 implementation, the MBD places its prior directly on the forward in time migration rate *m_ij_*, while the SC applies it to the backward in time migration rate (see also Methods). Effectively, this results in a broader prior distribution on in the SC compared to the MBD analysis (S3 Fig). By directly editing the BEAST xml, we placed the prior distribution on in the SC analysis. With these settings, the MBD and the SC show identical coverage statistics (S2 Fig). Furthermore, the SC obtains a 1.5 times narrower HPD width and 1.2 times smaller RMSE compared to the MBD.

In summary, the SC and the MBD models perform similarly well in the endemic setting, i.e. when the population size does not change.

### Migration rate inference in epidemic outbreaks

The epidemic scenario is characterised by a sudden increase in the number of infected with stochastic fluctuations over time. The MBD can account for the growing population while the SC implementation in MultiTypeTree cannot. We distinguish between low and high sampling, where sequences of ≈ 10% and ≈ 50% of the infected are collected, respectively.

Overall, the MBD performs better than the SC over all scenarios and settings. As an example, for low sampling (Fig 2, right column), the MBD recovers the migration rates significantly better (≥ 93%) than the SC (≤ 87%) across migration cases. When we place the SC migration rate prior on the forward migration rate as in the endemic scenerio (S2 Fig), the coverage increases from 70% to 83%. When more individuals are sampled, the performance of the MBD does not notably change (compare S4 Fig). The SC attains a higher coverage of *m_rb_* (80% compared to 70%) for fast migration.

Further, the RMSE of *m_rb_* decreases from ≈ 76 to 10 but it increases for *m_br_* from 35 to ≈ 130. Hence, a high sampling proportion does not change the results qualitatively.

In summary, the MBD performs better for epidemic diseases or generally, when the population size increases exponentially. Migration rate estimates under the SC show low coverage and high RMSE. We hypothesise that these inaccurate estimates are due to the SC’s assumption of a constant population size. In the following section, we will investigate how exactly the SC fits the data when population structure and size changes are present.

### The SC and exponential growth

To understand the source of the SC’s inaccurate estimates, we visually inspected the coloured trees it infers (example in S6 Fig, panel A). For fast migration, we observed that migration mostly flows one-way, e.g. only from red to blue in the exemplary tree. This is corroborated by the inferred migration rates (S6 Fig, panel B), which are negatively correlated (*τ* = −0.7, p-value< 0.001). Further, the root location is tied to the higher migration rate, e.g. right of the black line in S6 Fig, panel B, the higher migration rate *m_rb_* always coincides with the root location *r*. That means the SC infers a migration process along the tree that starts in either of both locations and then only allows migration into the other location. This strongly deviates from the true migration process (S6 Fig, panel A). Further, only the coalescence rate from the root location is constrained, e.g. λ_*r*_, right of the vertical line in panel D while the HPD interval of the other can span between [1 − 0.003].

We do not find those patterns in the MBD estimates (S6 Fig, panel C, E). Hence, we can exclude that signal for the above mentioned correlations was present in the simulated data. For slow migration, the anticorrelation within and between migration and coalescent rates becomes less pronounced in the SC analysis (see S8 Fig). Thus, given stronger signal for population structure the SC can better estimate the migration rates in the presence of an exponentially growing population (see also increased coverage in Fig 2).

In summary, given weak signal for population structure (fast migration), the SC infers a specific migration process along the tree that deviates strongly from the truth. If the population is more structured, this pattern becomes less pronounced.

### Root location inference

The root state provides information on the potential source of an epidemic. Here, we compare how well the models recover the root location based on trees from the endemic and epidemic scenario. We calculate the posterior probability for the root node being in location *r* and *b*. Note that 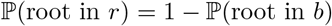. We assign the root a location if the posterior probability for either location is > 50% or > 90%. For the trees with a marked root location we report how often the true root location was assigned in Fig 3.

**Fig 3.**
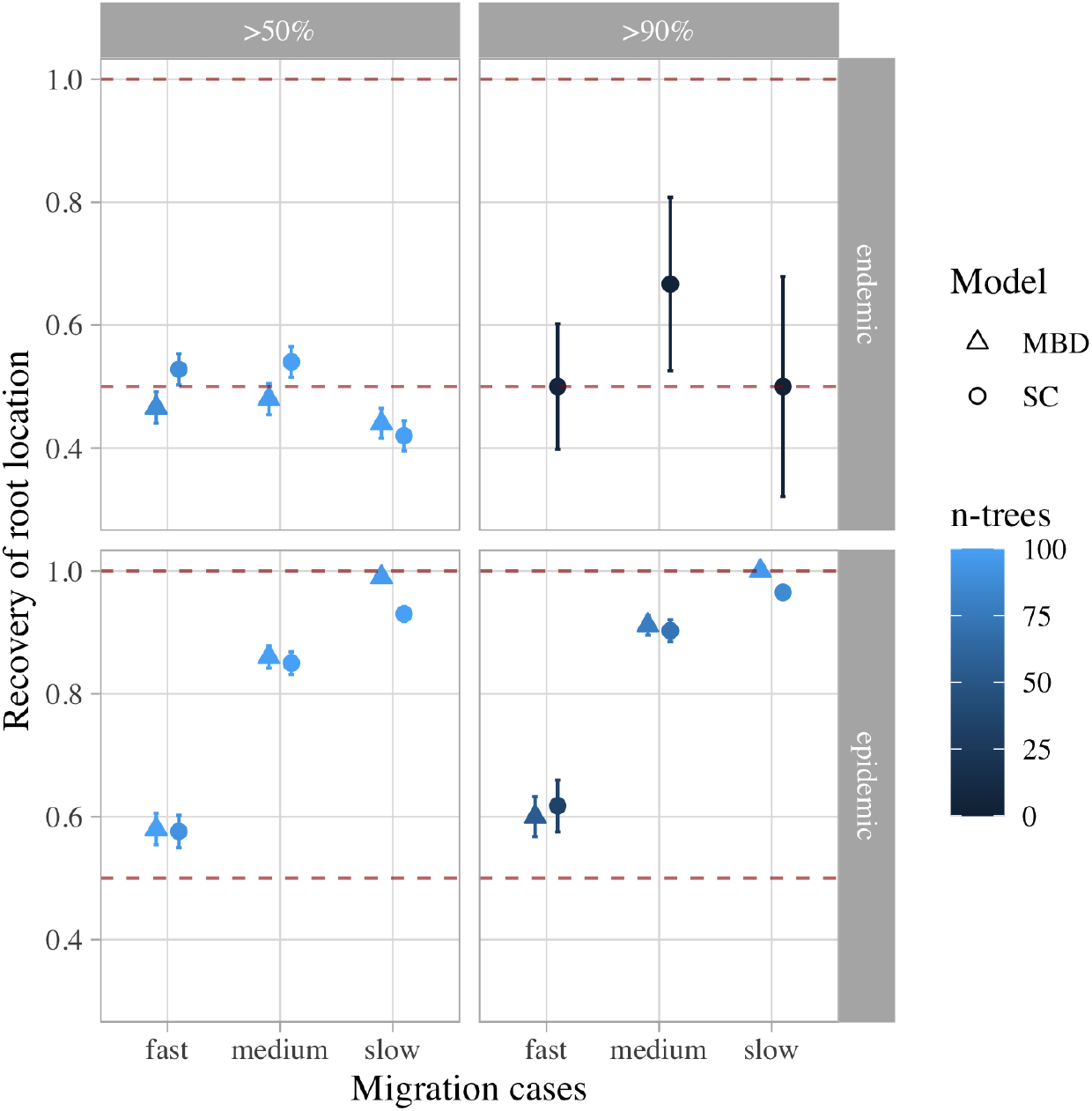
Inference of the root location. We show how the SC and the MBD recover the root location from endemic and epidemic trees. We vary the required level of certainty in the root assignment between > 50% and > 90%. The colouring displays how many trees meet the requirement. The recovery is displayed for every migration case. The red dashed lines indicates the recovery for optimal (1.0) and random recovery (0.5).

For the endemic scenario, both models recover approximately half the root locations correctly. The MBD does not provide > 90% posterior support for the root state for any of the simulated trees. The SC does provide this level of support for 2 – 6 trees across migration cases. Its recovery fraction, however, only fluctuates around 0.5.

In the epidemic outbreak scenario, the models perform better at root state recovery. For the minimal posterior propability of > 50% and fast and medium migration, both methods recover the true root state similarly well. For slow migration, the MBD’s recovery (99%) is 5 percentage points higher than the SC’s. Increasing the minimum posterior requirement for root assignment improves the root recovery for fast and medium migration for both models. Dense sampling (see S7 Fig), mostly influences the SC’s results. It improves the recovery for slow (+5 percentage points) while worsening it for fast migration (−8 percentage points).

There are several reasons why the models recover the root state better from the epidemic trees. A major influence are the sampling times. In the epidemic scenario, we sample continuously through time resulting in samples close to the root (compare Fig 1). In contrast, the endemic trees only have samples close to the present. Hence, while for the epidemic tree a simple parsimony assignment of the root state will often be correct, for the endemic trees the lineage locations have to be correctly backtraced throughout the tree. Additionally, the endemic trees are on average longer than the epidemic trees, which, all other things being equal, is bound to increase the uncertainty in lineage location.

Overall, both models recover the root state similarly well. This finding is interesting, given that the SC estimates inaccurate migration rates from epidemic trees. It appears that the inference of the root location is more robust against model misspecifications.

## Discussion

We compared how well the structured coalescent (SC) and the multitype birth-death (MBD) models infer migration rates and the root location from phylogenetic trees. We tested their performance for different parameter regimes, inluding endemic versus epidemic disease, varying migration rates and sampling intensities.

For endemic diseases (constant population size), both methods recover the true migration rates similarly well. While the MBD shows a higher coverage for medium migration, the SC consistently obtains narrower posterior intervals of the migration rate. Our results are consistent with previous analyses of coalescent and birth-death models for unstructured populations by Boskova et al. [22]. There, coalescent models obtained narrower posterior intervals than the birth death models when estimating the growth rate from coalescent trees. As Boskova et al., we hypothesize that the birth death model’s assumption of stochastic population size changes leads to wider posterior intervals as a wider class of parameter combinations are compatible with the observed tree.

For epidemic outbreaks (exponentially growing populations), the MBD infers migration rates with higher coverage and lower RMSE than the SC across parameter regimes. The traditional, constant-rate SC obtains these inaccurate estimates, because it cannot account for exponential growth. We show, that it infers a particular kind of migration process that deviates from the truth when signal for population structure is weak. Our findings motivate further research to combine the exponential growth coalescent with population structure.

For both the endemic and epidemic scenario, we found that it matters for migration rate inference whether the SC prior is placed on the forward or backward migration rate. Hence, practitioners should consider whether their prior knowledge is best expressed in terms of the emigration (forward migration) or the immigration (backward migration) rate. We implemented an option to place the prior on the forward migration rate in SC analyses in the MTT package.

Both models infer the root location with similar coverage throughout parameter regimes. This is interesting because the SC estimates inaccurate migration rates in the epidemic scenario. Therefore, our simulations reveal that inferring the root state is more robust compared to the migration rates.

We show here that phylodynamic models can reliably extract information about population structure from phylogenetic trees. Such phylodynamic analyses complement confirmed case data analyses, which by themselves reveal little information about the underlying population structure. We envision that a future framework allowing for a simultaneous, coherent analysis of both data types will provide rapid and reliable information on epidemic spread in structured populations which is extremely valuable information for authorities to design public health interventions.

## Materials and methods

### Simulations

We simulated phylogenetic trees, that capture important characteristics of an endemic disease and an epidemic outbreak. An outbreak is characterised by an exponential increase in the number of cases of a disease. When case counts are still low, they vary in a stochastic way. The outbreak can be adequately modelled by the multitype birth-death (MBD) model, as it can capture exponential growth and explicitly accounts for stochastic population size changes. In an endemic situation, a pathogen is constantly present within a population. Without sudden changes in the number of infected, the structured coalescent (SC) model is an adequate model to simulate trees for that scenario.

For both scenarios, we assume two geographic compartments, which we label red (r) and blue (b). Within the compartments, individuals can transmit the disease to each other at random. Between the compartments *i* and *j*, individuals can migrate at rate *m_ij_*, with *i,j* ∈ {*r, b*}. We investigate three migration scenarios. Firstly, an infected individual is expected to change locations once in its lifetime (fast scenario). In the two slower migration scenarios, only one individual in 10 and 100 (medium and slow scenario) is expected to relocate. We simulate trees under these model settings and condition on each tree to have 50 tips from each location. That way, we ensure a similar amount of input information. In what follows, we provide further details on the simulation setup for the epidemic and endemic settings.

#### Epidemic simulations

For each compartment *i*, the MBD is parameterised by per-individual birth *β_i_* and death *δ_i_* rates and a sampling proportion *s_i_*. Additionally, the migration rate *m_ij_* parameterises the expected rate at which individuals in compartment *i* migrate to compartment *j*. In the epidemiological context, the birth rate is the rate at which an infected individual transmits the disease to an uninfected individual. The death rate is the rate at to which an infected individual becomes non-infectious, e.g. by recovering from the disease or quarantining. Upon removal from the pool of infected, an individual is sampled with probability *s_i_*, which is the proportion of sampled individuals in *i* from the pool of recovered individuals in *i*.

MASTER [23] was used to generate MBD trees for varying parameters. We set the birth rate *β_i_* = 1.5 and the death rate *δ_i_* = 1 for both locations. Hence, the epidemic grows exponentially at rate *r_i_* = *β_i_* − *δ_i_* = 0.5 in expectation. The migration rates *m_ij_* == *m_ji_* from fast to slow are 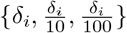. The simulation begins with one individual at time *t* = 0 in location *i* sampled uniformly at random from {*r, b*}. Trees are grown until the number of infected irrespective of region reaches a defined threshold *N_f_*. Then, 50 samples are drawn from all leaves of each location in the tree. As a leaf in a tree corresponds to a death event, we can approximate the sampling proportion for location *i* as follows:

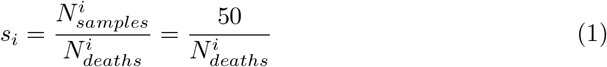

We choose two values of *N_f_*, such that the approximate sampling proportion corresponds to low sampling (*N_f_* = 300, *s_i_* ~ 0.1) or high sampling (*N_f_* = 100, *s_i_* ~ 0.5). The parameter combinations are summarised in Tab 1. For each parameter set, we simulate 100 trees.

**Table 1.** Parameter combinations for simulating MBD trees. Columns name the variables of interest. *β_i_* = 1.5, *δ_i_* = 1.0 for *i* ∈ {*r, b*}.

| Case | sampling | migration |
| --- | --- | --- |
| 1 | low | fast |
| 2 | low | medium |
| 3 | low | slow |
| 4 | high | fast |
| 5 | high | medium |
| 6 | high | slow |

#### Endemic simulations

The SC with two compartments *i* and *j* is parameterised by the pairwise coalescence rates λ_*i*_, λ_*j*_ and the backward migration rates *q_ij_, q_ji_*. The SC parameters quantify processes that occur backward in time, i.e. from the tips of the tree toward the root. The coalescence rate is the rate at which a pair of lineages coalesces as we move from the tips of the tree toward the root. The migration rate *q_ij_* is the rate at which an individual from location *i* moves to location *j* backward in time. We start the simulation at the leaves of the tree in the present and merge the lineages together as we go into the past. For each tree, we draw 50 leaf times for each location uniformly at random between (0,10). For the MBD we defined the migration rates as fast, medium and slow with respect to the birth rate. For the SC we use [24] to determine the coalescence rate which corresponds to the birth rate and the final population size employed under the MBD model, which we used in the epidemic simulations:

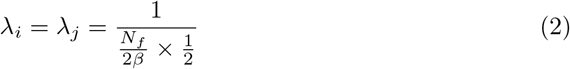

The factor 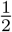 arises because we assume that the final population size *N_f_* consists of individuals from either location in equal shares. Hence, we set 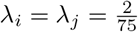. Then, we convert the forward migration rates from the epidemic simulation into the backward migration rates [25]:

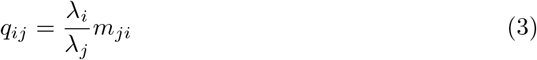

Hence, we set *q_ij_* to 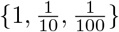 for fast, medium and slow migration. Then, we use MASTER [23] to simulate 100 SC trees for every migration case as before.

### Inferences

We fit the models with packages implemented in the BEAST 2 [26] Bayesian inference framework. We use the BDMM [5] package for the MBD and the MultiTypeTree [3] package for the SC inference. In the MBD analysis, we infer the posterior for three parameters for each location i: the birth rate *β_i_*, the sampling proportion *ψ_i_* and the forward migration rate *m_ij_*. We fix the death rates in both locations *δ_r_* = *δ_b_* = 1 to avoid unidentifiability constraints.

In the SC inference, we estimate the posterior distribution of two parameters for each location i: the coalescence rate λ_i_ and the backward migration rate *q_ij_*. To compare the inferred migration rates between both models, we convert the SC’s backward migration rates *q_ji_* to forward migration rates *m_ij_* by rearranging Eq. 3.

In both sets of analyses, we fix the tree to the truth, because we are only interested in the differences arising from our tree prior choice (MBD or SC). The prior distributions on all parameters can be found in S1 Table and S2 Table. All MCMC analyses were run for 10^8^ steps and 10% of burn-in was discarded. Additionally, we verified that the effective sample size was higher than 200 for all parameters, to ensure the runs were converged.

#### Summary statistics

To compare the posterior distributions on *m_ij_* inferred by the SC and the MBD, we chose the following three metrics: coverage, relative 95% highest posterior density (HPD) width and relative root mean square error (RMSE). The coverage quantifies, how often the true parameter is contained in the credible interval of the posterior. Here, we use the 95% HPD, which is the narrowest region of the posterior, that holds 95% of the probability mass. The second metric, HPD width, is the width of that interval and measures the precision of the estimate. Lastly, we compute the RMSE between the inferred median and the true value, which quantifies how accurate the estimate is. Both the HPD width and the RMSE are divided by the true migration rate and hence we obtain their relative versions. Therefore we can compare them across the different migration cases.

For each model and parameter case, we have 100 independent MCMC chains (one for each tree). For each parameter of interest we compute the posterior median and boundaries of the posterior 95% HPD interval from a MCMC chain (for a given tree). Given this summary data, for each of the previously mentioned metrics we compute the mean and standard deviation by bootstrapping over 1000 replicates using the R Boot package [27]. We use bootstrapping because the distribution of the posterior medians is not necessarily normal.

#### Root Location

For each tree, we compute the posterior probability of the root being in location *r* and *b*. Note that 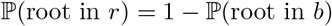. We consider two methods for assigning the root state. First, if the posterior probability for root state *r* is above 50%, we assign the root the state *r*, and otherwise b. Second, only if the posterior probability for a root state is > 90%, we assign that particular root state; otherwise we assume that the root state cannot be determined. For every scenario and parameter case we compute the recovery, defined as the number of times the root was assigned to the correct location. We use bootstrapping to determine the mean and standard deviation or the recovery as before.

## Acknowledgements

The authors thank ETH Zurich for funding.

## Author Contributions

Conceived and designed the simulation study: SS TS TV. Performed the simulations: SS. Analyzed the data: SS TS TV. Contributed reagents/materials/analysis tools: SS TS TV. Wrote the paper: SS TS TV.

## Data Availability

All code to rerun the analysis and generate the figures is available from https://github.com/seidels/structured-phylodynamic-models.

## Supporting information

**Fig S1.**
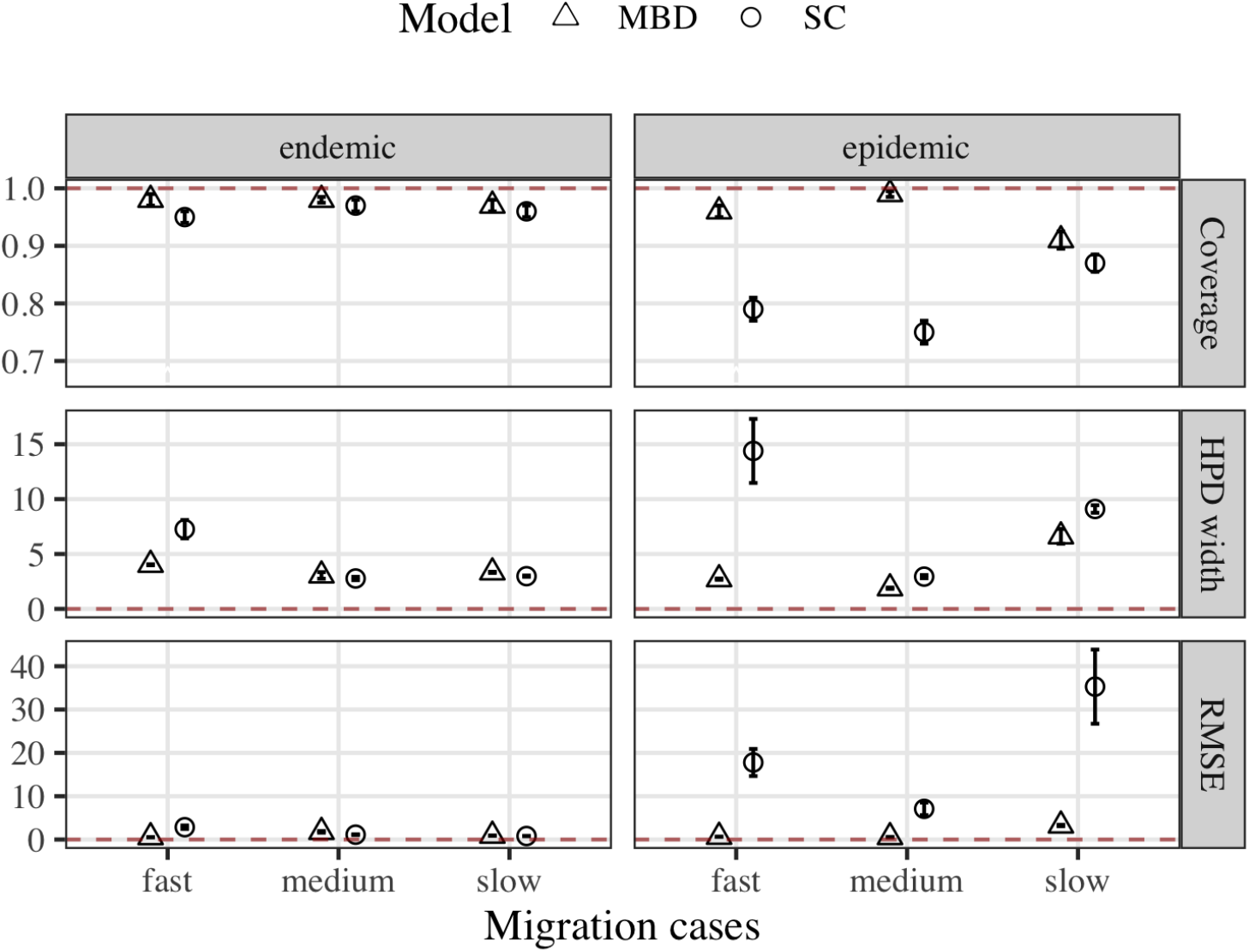
Summary statistics for *m_br_*. For legend see Fig. 2

**Fig S2.**
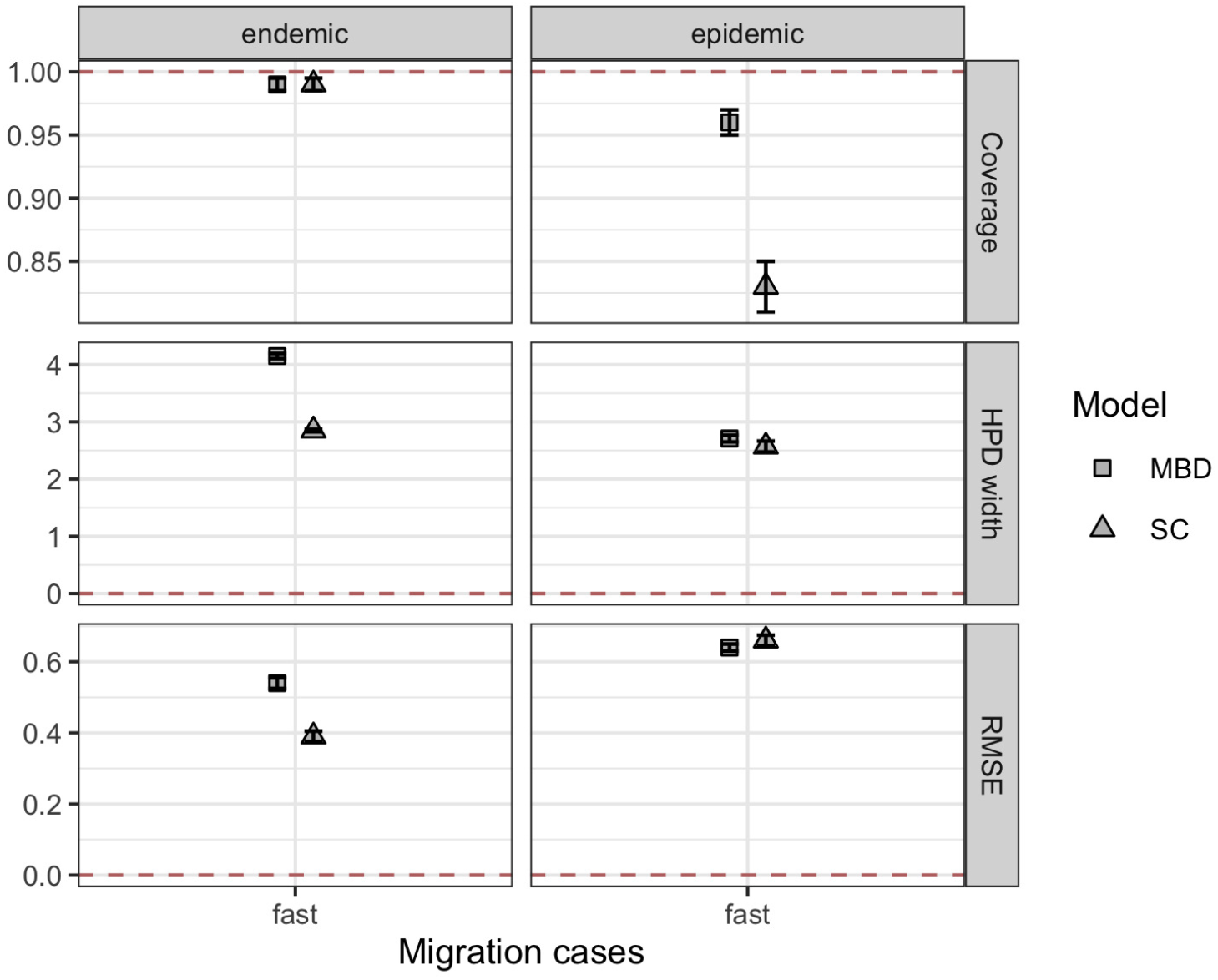
Summary statistics for fast migration and identical priors. Here, we show the summary statistics for *m_rb_* in the endemic and epidemic scenario, when identical priors are placed on the forward migration rate *m_rb_* in the SC and MBD analyses. For legend refer to Fig. 2

**Fig S3.**
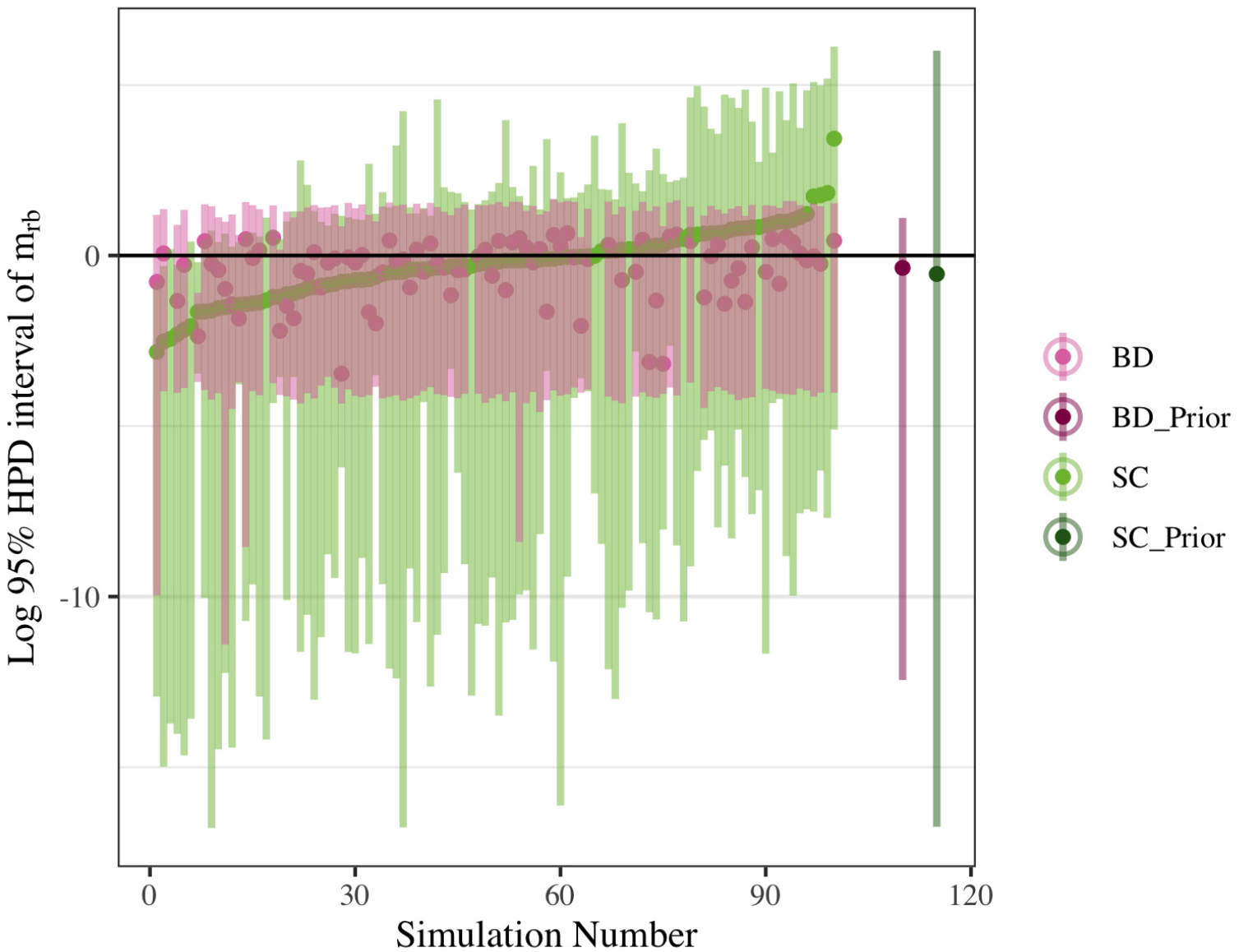
Influence of the prior for endemic disease. For every simulated tree (Simulation number) in the endemic scenario with fast migration, we plot the inferred 95% HPD intervals as lines and medians as dots. Estimates are shown in light purple for the MBD and in light green for the SC analysis. The black line indicates the true migration rate. On the right, the MBD’s prior distribution on mrb is shown in dark purple and the SC’s counterpart in dark green. Placing the SC’s prior on the backward migration rate *q_br_* effectively broadens the prior distribution over *m_rb_* in comparison to the MBD. In the inference, the MBD’s prior seems to restrict the posterior from higher migration values.

**Fig S4.**
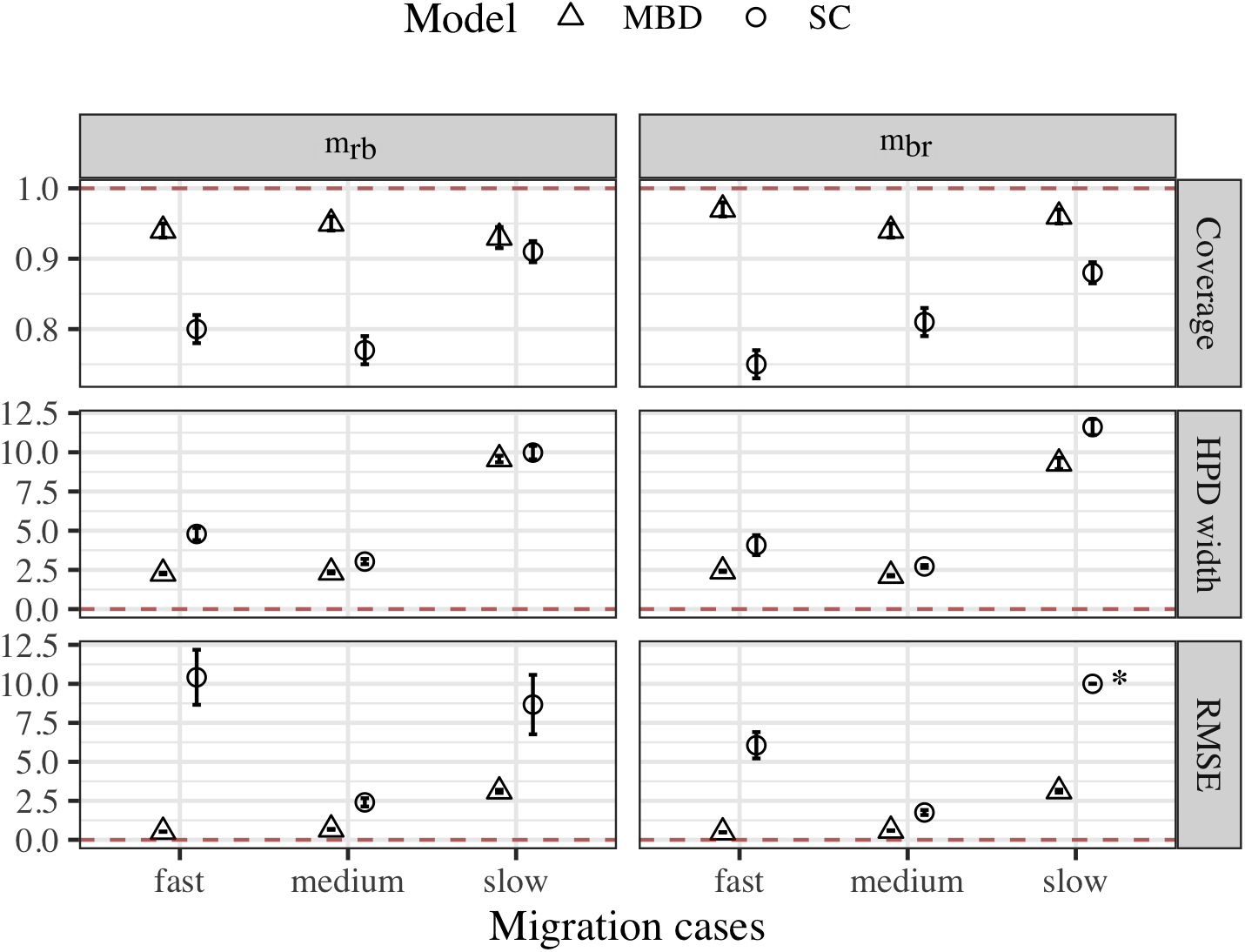
Summary statistics for epidemic scenario and high sampling. Results for *m_rb_* and *m_br_*. For legend see Fig. 2. The SC RMSE for *m_br_* and slow migration was set to 10 for better visibility of the differences between the other RMSE values. Its true value is 130.5 ± 94.

**Fig S5.**
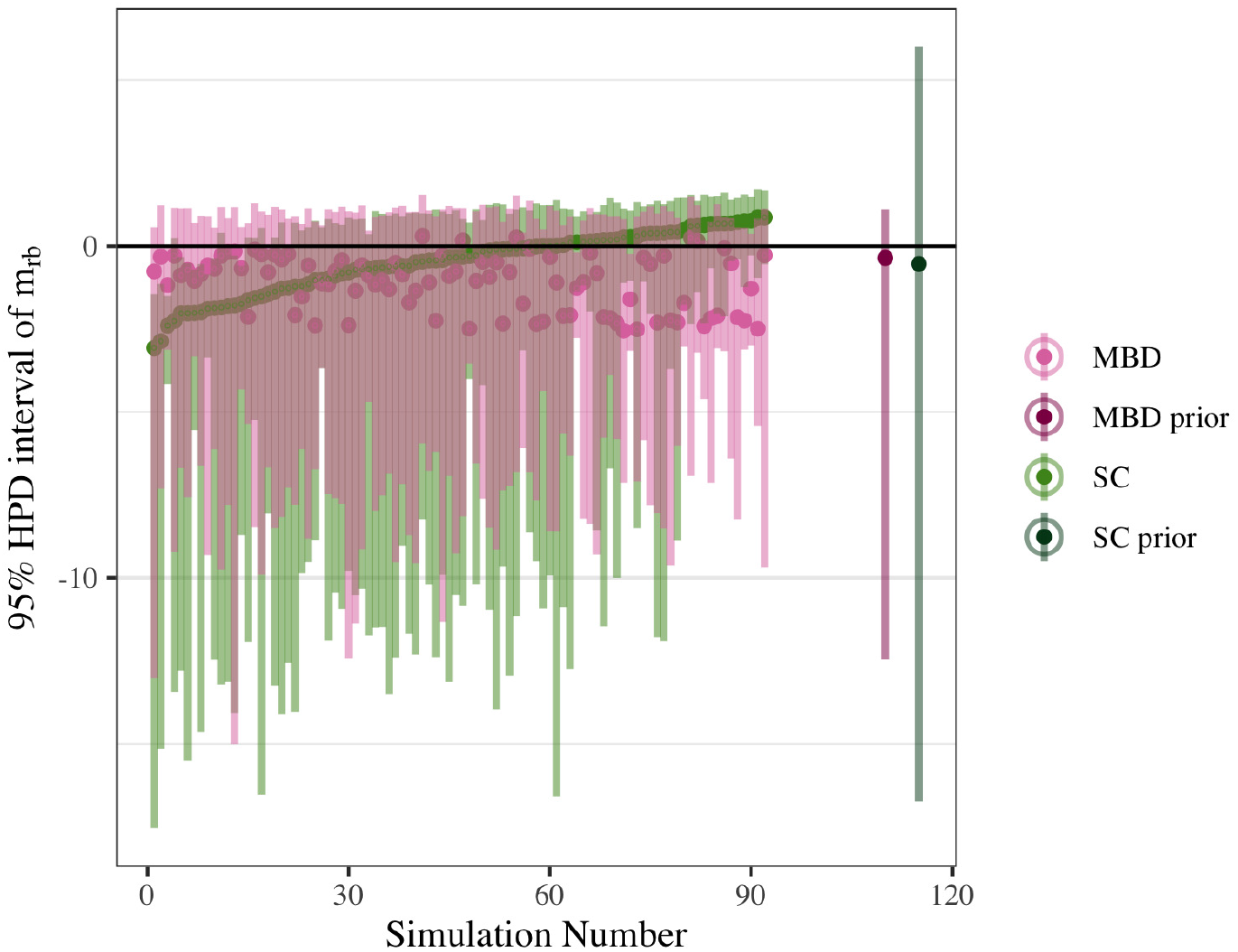
Posterior HPD intervals and influence of the prior. HPD intervals and medians of *m_rb_* are shown for the epidemic scenario with fast migration. Refer to legend in S3 Fig

**Fig S6.**
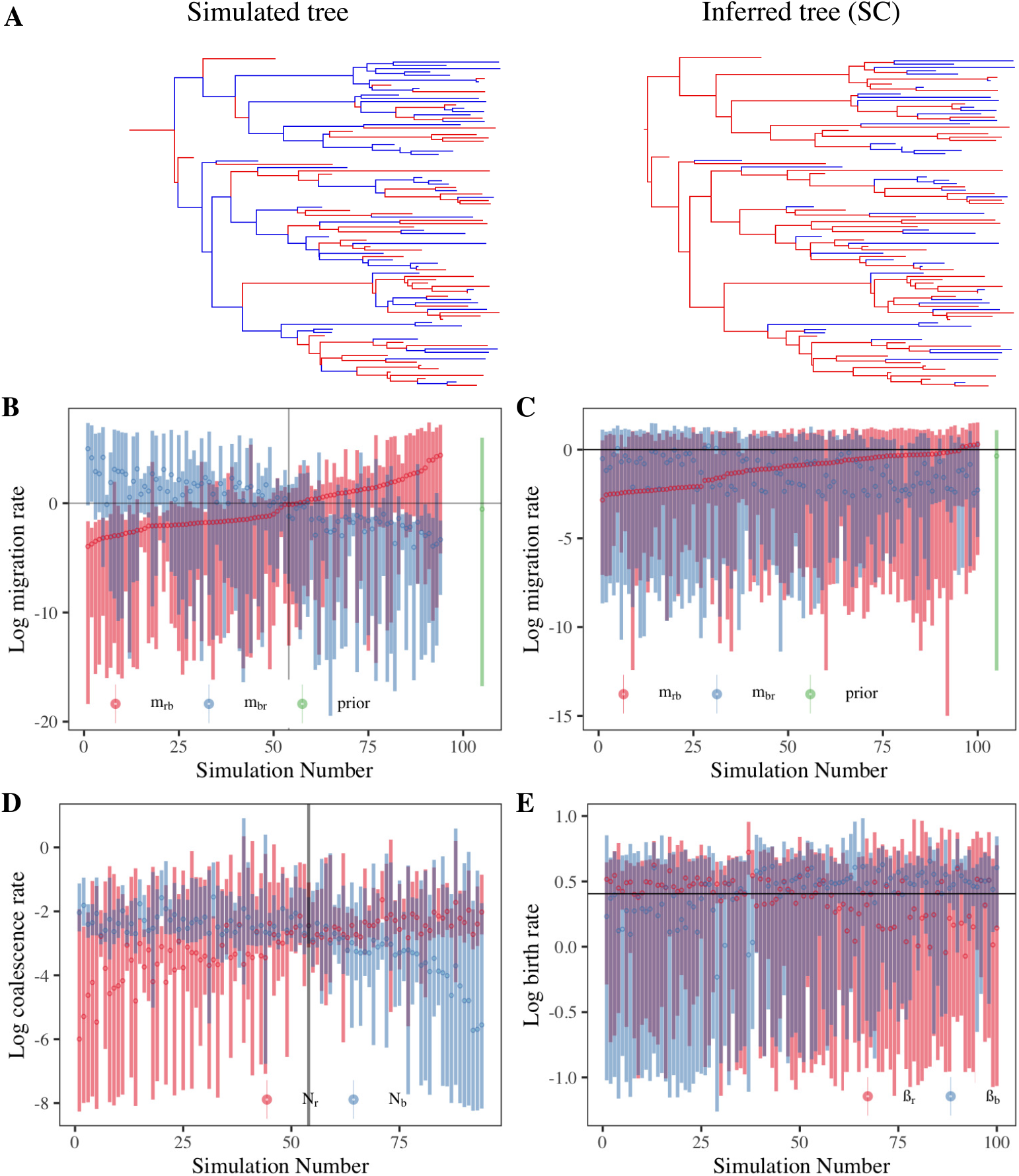
SC analysis of exponential growth trees. The exemplary trees show a tree simulated under exponential growth and fast migration (left) and the inferred lineage locations (for color legend see Fig. 1) by the SC. The SC recovers a tree with notably less migration events than in the simulation. (**B, C**) The log-transformed, median migration rates *m_ij_* (dots) and the HPD intervals (shaded regions) inferred by the SC (**B**) and the MBD (**C**) are plotted for each simulation number (tree). The simulation numbers are sorted, s.t. mrb is increasing. The black, horizontal line indicates the true migration rate. On the left of the vertical line the inferred root location is b and vice versa. For the MBD, the inferred root locations are not separable by a vertical line. (**D, E**) The median coalescence rates *N_i_* (SC, **D**) and birth rates β_i_ (MBD, **E**) and their HPDs are plotted against each simulation number. The ordering of the simulation numbers is taken over from the respective upper panel.

**Fig S7.**
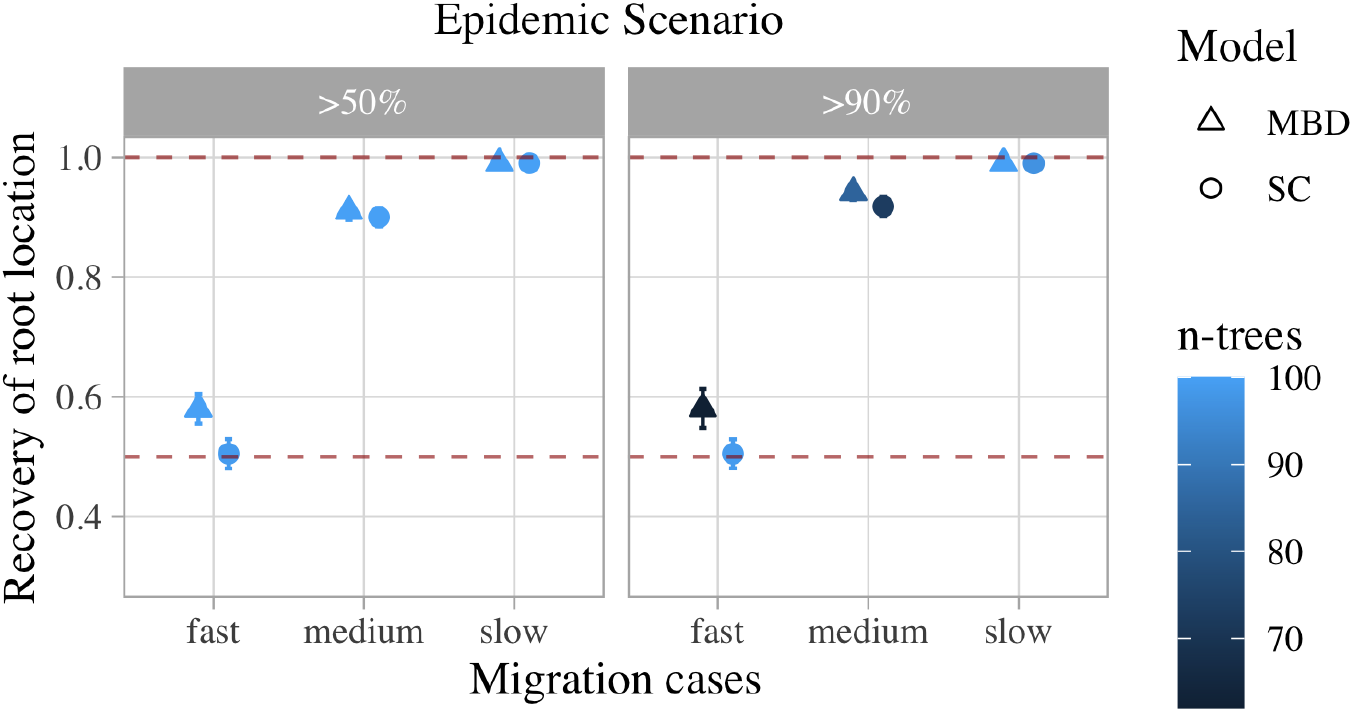
Root state recovery for epidemic scenario with high sampling. For legend, see Fig. 3

**Fig S8.**
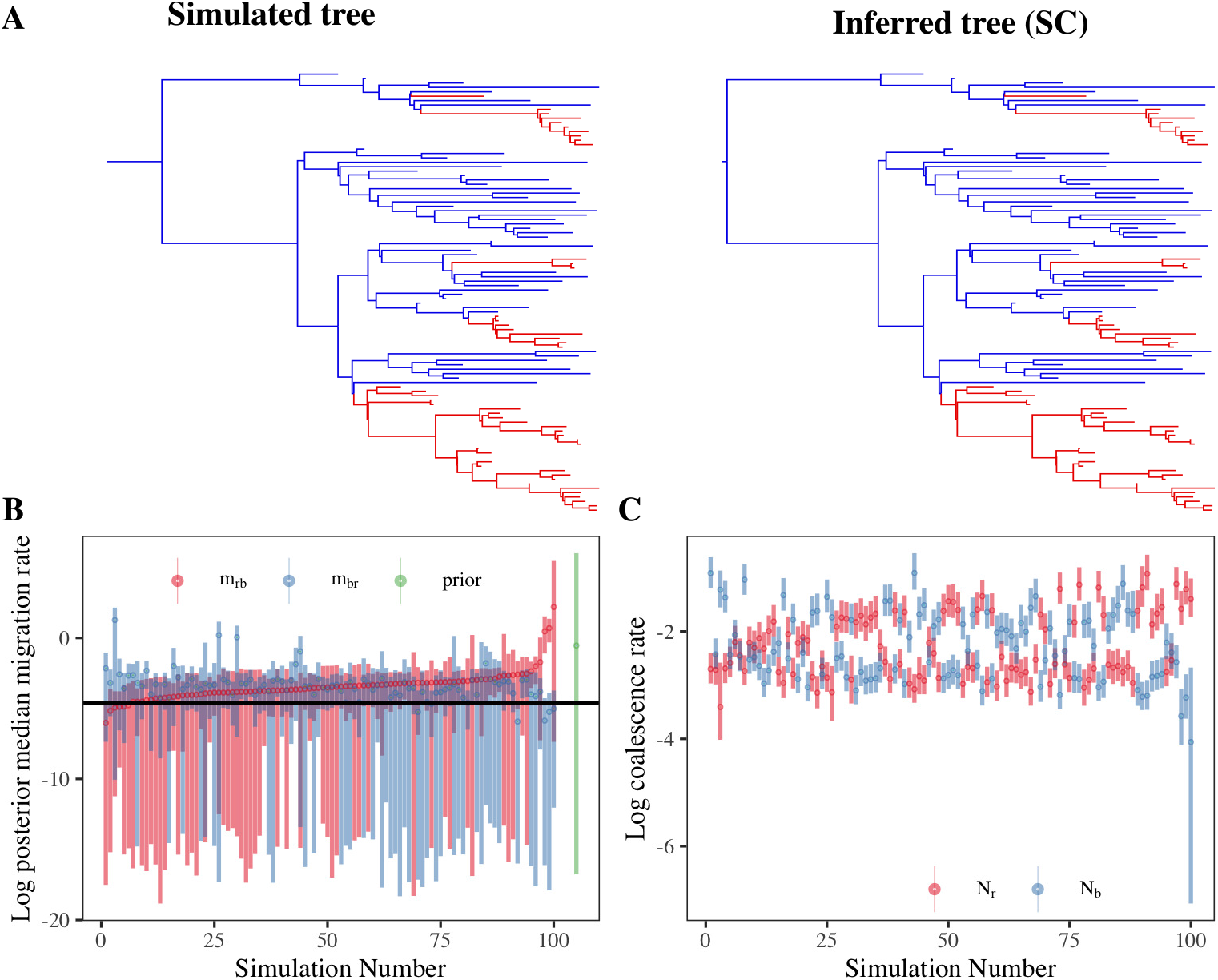
SC inference from exponential growth trees for slow migration. Find the legend in Fig S6 Fig. Note that panel C in this figure corresponds to panel D in Fig. S6 Fig.

**Table S1.**
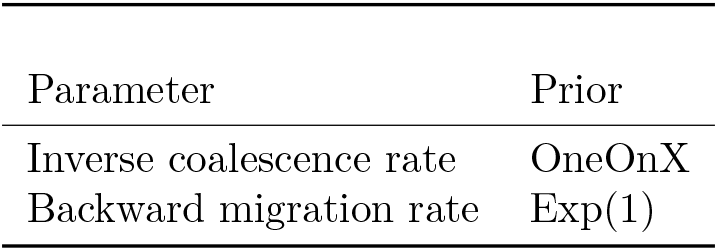
Priors on parameters in structured coalescence inference.

**Table S2.** Priors on parameters in multitype birth-death inference.

| Parameter | Prior |
| --- | --- |
| Birth rate | LogNormal(0.25, 0.5) |
| Forward migration rate | Exp(1) |
| Sampling Proportion | Uniform(0,1) |

